# Distinct tau filament folds in familial frontotemporal dementia due to the *MAPT* S305I mutation

**DOI:** 10.64898/2026.02.12.705620

**Authors:** Henry S. Pan, Gregory E. Merz, Alissa Nana Li, Minh Quan Le, Hyunil Jo, Athena Quddus, Anthony Yung, Rian C. Kormos, Arthur A. Melo, Eliana Marisa Ramos, Argentina Lario Lago, Salvatore Spina, Lea T. Grinberg, Howard J. Rosen, Eric Tse, Maria Luisa Gorno-Tempini, William F. DeGrado, William W. Seeley, Daniel R. Southworth

## Abstract

Frontotemporal lobar degeneration with tau inclusions (FTLD-tau) comprise a class of fatal heterogeneous neurodegenerative diseases. Approximately 10% arise from pathogenic MAPT mutations and often cause severe, early-onset disease with pathology that is distinct yet partially overlapping with sporadic cases. Here, we evaluated post-mortem tissue from a patient with FTLD-tau due to *MAPT* S305I showing neuropathology most consistent with argyrophilic grain disease (AGD), a prevalent limbic tauopathy of aging. Structures determined by cryo-electron microscopy reveal tau filament folds that differ from those found in sporadic AGD or other tauopathies and feature a 4-layer architecture stabilized by the Ile substitution within its core. Comparative structural analysis reveals conserved motifs are shared among AGD, corticobasal degeneration, and *MAPT* P301T. A well-defined density stacks along a cationic cleft, indicative of a bound RNA-like polyanion or small-molecule. *In vitro* analysis shows the S305I mutation promotes fibrilization relative to normal tau. These results demonstrate that *MAPT S305I* stabilizes a distinct aggregation-prone tau fold that likely contributes to disease pathology and heterogeneity beyond its known splicing defects, and underscore potential limitations of using the most pathologically similar genetic form as a model for sporadic FTLD-tau.

## Introduction

Tauopathies, including Alzheimer’s disease (AD), progressive supranuclear palsy (PSP), corticobasal degeneration (CBD) and others, are a diverse group of neurodegenerative diseases characterized by the accumulation of toxic protein deposits in the brain comprised primarily of the microtubule-associated protein tau (MAPT, or tau)^1–9^. Cryo-electron microscopy (cryo-EM) studies of tau filaments extracted from post-mortem patient brain tissue have revealed that the structural core, which typically includes microtubule-binding repeat domains (R1-R4) and a portion of the C-terminus (C) adopts different cross-β amyloid folds in disease^10–17^. These insights have advanced disease classification by identifying specific types of tau filament folds that arise in different pathologically defined tauopathies^10^. Moreover, the diversity of tau filament structures indicates different molecular mechanisms likely underlie tau aggregation in these diseases. Nonetheless, the molecular and pathological factors that drive the formation of a particular tau disease fold remain poorly understood.

Investigations of pathogenic *MAPT* variants have established the central role of tau aggregation in disease^18^ and offer important insight into the molecular drivers of tau filament folds. While most tauopathies occur sporadically^19^, more than 60 familial mutations in *MAPT* have been identified to date, each associated with a mutation-specific pattern of frontotemporal lobar degeneration with tau pathology (FTLD-tau)^20^. The observed FTLD-tau pathomorphological features overlap with those seen in the sporadic forms, including AD, PSP, CBD, Pick’s disease (PiD), and argyrophilic grain disease (AGD)^10,21–23^, prompting some authors to propose that sporadic FTLD-tau can be effectively modeled using familial *MAPT* mutations^24^. Mutations at residue P301, including P301L and P301S, are known to enhance tau aggregation and seeding and are widely used in cell and transgenic animal models for tauopathies, including AD^25–29^. Cryo-EM studies have revealed that familial tau mutations form specific filament folds: P301L and P301T adopt unique structures^30^, while V337M and R406W are similar to the AD paired helical filament (PHF) fold^14^. However, only a limited number of tau structures from patients with *MAPT* mutations have been characterized, despite strong interest in defining tau folds that arise in disease and advancing the development of small-molecule diagnostics and therapeutics that selectively bind specific folds^31–33^.

Familial mutations in *MAPT* occur in both exonic and intronic regions, including several at the exon 10 splice junction, which contains a stem loop regulatory element required for alternative splicing of 3-repeat (3R) and 4-repeat (4R) tau isoforms^34^ (**Fig. 1a**). Mutations in this region cause splicing defects that increase 4R tau levels^35,36^ and are proposed to drive 4R-predominant tauopathies resembling CBD and PSP. The *S305I* mutation, located within the coding sequence at the exon 10 splice junction, was previously identified in a patient with severe early onset FTD, with symptoms developing at age 39 and a rapid, 1.5-year disease course^37^. Post-mortem analysis revealed neuropathology similar to sporadic AGD, with argyrophilic grains, neuronal cytoplasmic inclusions, and oligodendroglial coiled bodies comprised of 4R tau, while classical neurofibrillary tangles, Pick bodies, and other tau morphologies were absent^37^. Sporadic AGD is the second most common tauopathy after AD^38^, and it is characterized by a very late age of onset and pathology that is typically limited to the limbic system, in contrast to the tauopathy resulting from *S305I* in which tau aggregation is more widespread in frontal and temporal regions,^37^ thereby producing an FTD-like clinical picture. Because inherited forms of FTLD-tau have been proposed as models for sporadic FTLD-tau subtypes^24^, it is important to determine whether the inherited and sporadic forms are associated with the same underlying tau filament fold.

**Figure 1:**
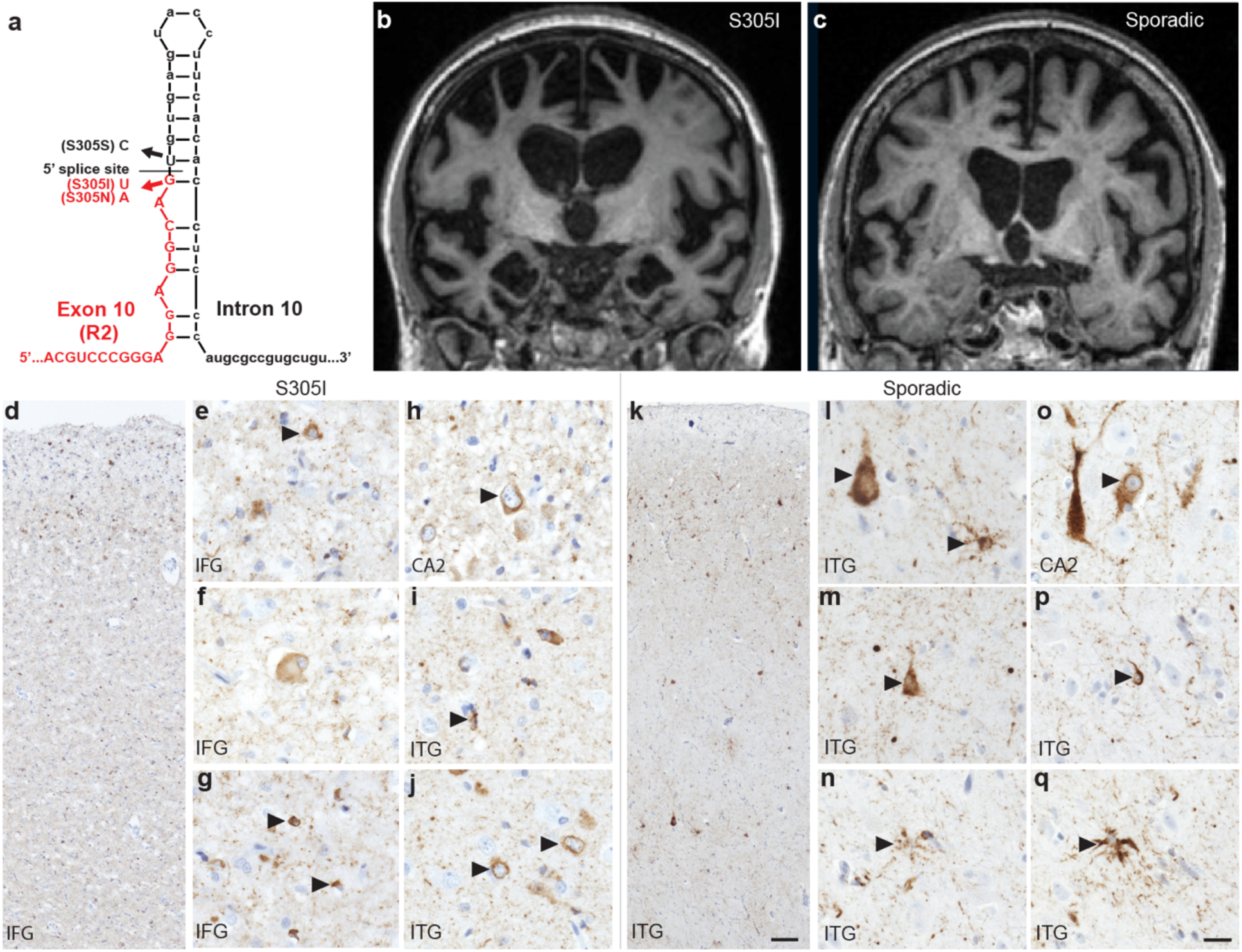
The *MAPT S305I* mutation and neuropathology. **a,** Schematic showing the location of *S305I* (with *S305S* and *S305N*) in the *MAPT* gene at the exon 10 (red) and intron 10 (black) splice-site junction and stem loop region. **b, c,** MRI scans from the patient carrying the S305I mutation (**b**) and a patient with sporadic AGD (**c**), shown for comparison. **d-q,** Representative images of immunostaining for phospho-tau (S202, CP13 antibody) in the patient carrying an S305I mutation **d-j,** and the patient with sporadic AGD **k-q**. Features of AGD are indicated by arrowheads in the inferior frontal gyrus (IFG, **d-g**), the CA2 region of the hippocampus (**h, o**), and the inferior temporal gyrus (ITG, **i-n, p, q**). Shared features characteristic of AGD include diffuse/granular cytoplasmic inclusions with perinuclear halos (**e, h, j, l, m, o**), and coiled bodies (**g, i, p**). Distinguishing features include ballooned neurons in the S305I case (**f**) and granular/fuzzy astrocytic inclusions in sporadic AGD (**l, n, q**). Scale bar in **k** = 100 µm and applies to **d** and **k**, and scale bar in **q** = 20 µm and applies to **e-j** and **l-q**.

Here, we sought to characterize the neuropathology and tau filament structures from a 39-year-old patient with the *MAPT S305I* mutation. Histopathological analysis revealed tau aggregates with morphological similarities to sporadic AGD but with more severe pathology, extending into the frontal and temporal cortices. Cryo-EM structures of tau filaments isolated from post-mortem tissue were determined to 3.1 and 3.2 Å resolution and demonstrate that S305I tau adopts two unique single protofilament folds. Each is comprised of an identical 4-layer fold and β hairpin involving microtubule repeat domains R2-R4, as well as an R1:C arm arranged differently in the two structures. The S305I residue contributes to the overall fold by packing in a solvent-inaccessible hydrophobic pocket in the center of the 4-layer core. Notably, a distinct unassigned density is resolved along a solvent-facing cleft formed by basic residues in R2 and R3 that is suggestive of a cellular polyanion, such as RNA, or a stacking anionic metabolite. Finally, *in vitro* fibrilization analysis of recombinant tau protein containing the S305I mutation reveals more rapid poly-phosphate-induced aggregation relative to normal tau, indicating the mutation alone contributes to tau aggregation. These results reveal that the *MAPT* S305I mutation results in a distinct 4R tau filament fold not previously observed in other sporadic or familial tauopathies. Moreover, S305I may contribute to the unique early-onset AGD-like pathology through accelerated aggregation and promotion of a specific tau filament fold, in addition to previously reported splicing defects and increased 4R tau levels.

## Results

### *MAPT S305I* FTLD-tau Case with Distinct and AGD-like Neuropathological Features

Here, we studied a 39-year-old woman who developed subtle changes in social behavior at age 35, followed by difficulties with speech, spelling, and grammar, prompting a diagnosis of nonfluent variant primary progressive aphasia. Later, she developed more widespread cognitive deficits, including memory loss. Brain MRI revealed severe grey and white matter atrophy most notable in dorsal and opercular frontal, medial temporal, insular, inferior parietal, and limbic structures (**Fig. 1b**). These findings are unlike those typically associated with sporadic AGD, in that less severe atrophy and more focal limbic atrophy is seen (**Fig. 1c**). Her father was diagnosed with FTD in his 50s. *MAPT S305I* was uncovered via research-based genetic screening that included *MAPT*, *APP, C9ORF72, FUS, GRN, MAPT, PSEN1, PSEN2* and *TARDBP* genes, as described previously^39,40^. Autopsy was performed ∼13 hours after death, and the fresh brain weighed 1000 grams. Hematoxylin and eosin staining of post-mortem brain tissue revealed severe degenerative changes in the frontal operculum, dorsal frontal regions, and angular gyrus that were more pronounced than typically seen in AGD^37^, as well as mild to moderate changes in ventral frontal and temporal areas. Immunohistochemistry for hyperphosphorylated tau (CP13 antibody) revealed a pattern of fine argyrophilic grains and threads in both cortical and subcortical regions, along with diffuse/granular neuronal cytoplasmic inclusions, at times with perinuclear halos, occasional compact round neuronal cytoplasmic inclusions, and coiled bodies in areas exhibiting degenerative changes (**Fig. 1d-j**). These features are similar but distinct from those seen in sporadic AGD (**Fig. 1l-q**). Granular/fuzzy astrocytes, a common though not universal feature of sporadic AGD, were absent^41^. An antibody against 3-repeat tau (RD3) disclosed normal findings, confirming the 4-repeat nature of the tauopathy. While these morphological findings are consistent with the previous characterization of a patient with the same *MAPT S305I* mutation^37^, we conclude that overall, *MAPT S305I* results in certain distinct neuropathological features compared to AGD.

### S305I Tau Filament Structures Adopt a Unique 4-Layer Fold

We next sought to characterize and determine structures of tau filaments isolated from the patient with *MAPT S305I* described above using cryo-EM. Sarkosyl-insoluble filaments were purified from 1g of fresh-frozen inferior frontal gyrus (IFG) tissue, as previously described^10^ (see Methods), and verified by negative-stain and cryo-EM to contain predominantly tau filaments with a wavy morphology (**Fig. 2a and Extended Data Fig. 1a-b**). Two-dimensional (2D) class averages exhibit filaments with two distinct morphologies (hereafter referred to as S305I tau Type I and Type II), based on the twist and a crossover distance, measured to be ∼600 Å for Type I and ∼750 Å Type II (**Fig. 2b**), each with well-defined 4.8 Å β-sheet spacing (**Extended Data Fig. 1c**). Notably, the crossover distances are considerably shorter than for sporadic AGD tau Types I and II, which are reported to be 240 nm and 216 nm, respectively^10^. Following 3D classification and helical reconstruction, we determined structures of S305I tau Type I and Type II filaments at resolutions of 3.1 Å and 3.2 Å, respectively, revealing two distinct filament folds that had not been previously reported (**Fig. 2c, Extended Data Fig. 1d-f, and Table 1**). Type I constituted the majority (∼85%) of the dataset, whereas Type II filaments were less abundant (∼15%).

**Figure 2:**
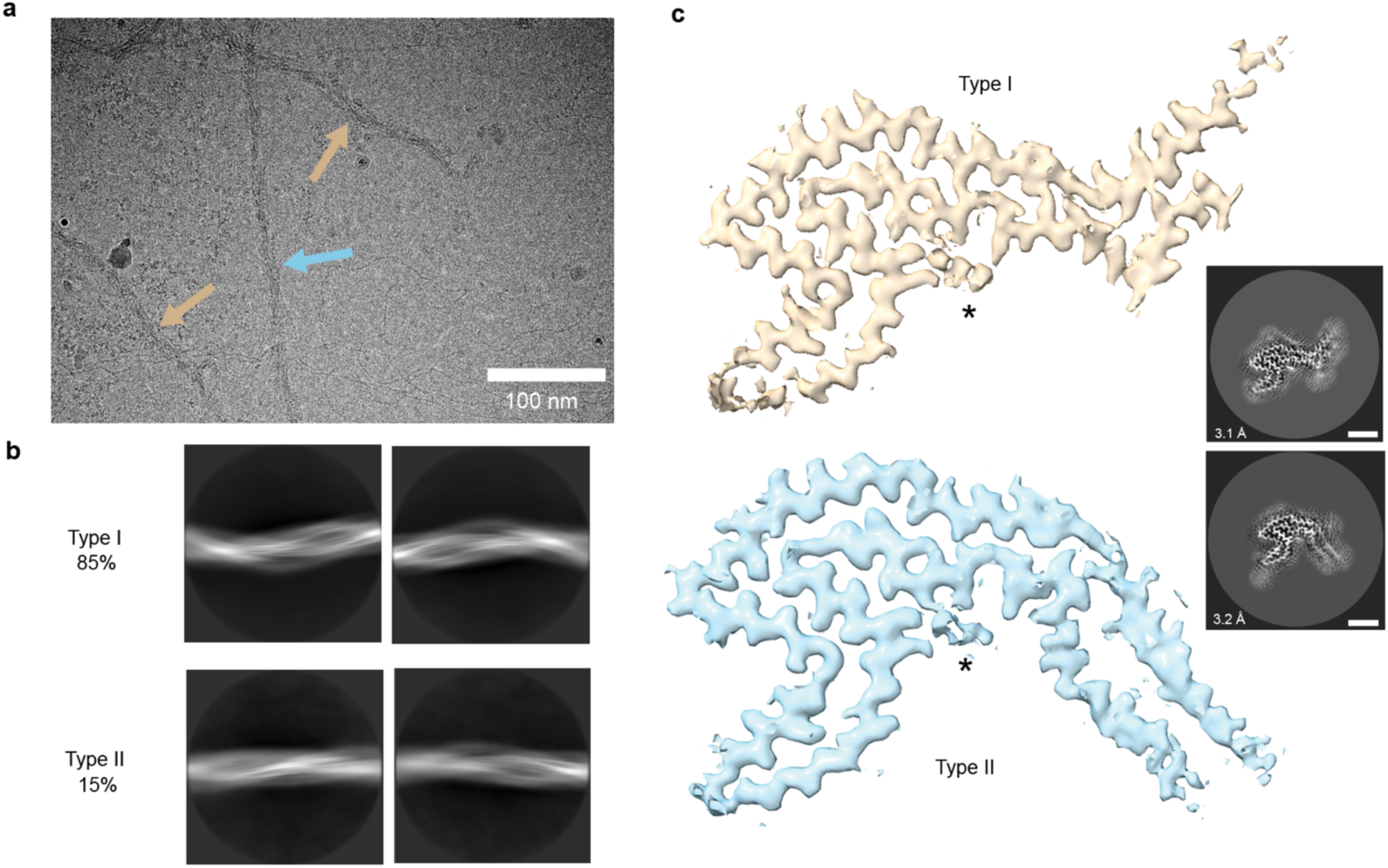
Cryo-EM 2D characterization and 3D structures identifying distinct S305I tau filament folds. **a,** Representative cryo-EM micrograph showing Type I (beige arrows) and Type II filaments (light-blue arrow). Scale bar equals 100 nm. **b,** Representative 2D class averages of Type I (top) and Type II (bottom) folds (with a 750 Å box size) showing distinct wavy morphologies. **c,** Cryo-EM final maps of S305I Type I (top, beige) and Type II (bottom, light-blue) tau filaments with additional, unaccounted density shown (*) and corresponding 2D cross-sections with resolution and particle percentages indicated for each final map shown (3.1 Å with 85% of the data for Type I and 3.2 Å with 15% for the data for Type II). Scale bars equal 30 Å.

The maps for both Type I and II filaments exhibit continuous, well-resolved density with side chain features, revealing both adopt a central 4-layered fold with an exposed hairpin on one face and a lower resolution stem at the opposite face that is positioned differently in each structure (**Fig. 2c**). Notably, a small but well-resolved discontinuous density is identified in a solvent exposed cleft of the 4-layer fold (discussed below). Human 4R tau residues G273–R379 were modeled into the map using Isolde^42^ based on a ModelAngelo^43^ predicted fit, revealing they form a large 97-residue folded hairpin established by distal cross-β contacts between R2 I277 and C I371 (**Fig. 3a-c**). The 4-layer core is comprised of R2-R4, with the entire length of R4 (residues 337-368) forming the solvent exposed face, contacting R2 (residues 275-285) and R3 (residues 305-307), which together form a 9-residue internal β-hairpin (**Fig. 3b-c**). Notably, I305 is well-resolved and located centrally in the 4-layer fold, pointed internally toward a 3-residue hydrophobic groove in R4 (discussed below). R3 turns away from the 4-layer fold, forming an 18-residue solvent-exposed β-hairpin (involving residues K317-G334). Alignment of the Type I and II structures reveals they are nearly identical at the 4-layer fold and β-hairpin regions (C_α_ root-mean-square deviation (RMSD) = 1.87 Å for residues K317-G334). However, the C-terminal arm positions of Type I and Type II filaments are substantially rotated (∼100 deg) relative to each other due to a sharp kink in Type I involving cross-β interactions D283:K369, S285:K369 and S285:G367 (**Fig. 3d**).

**Figure 3:**
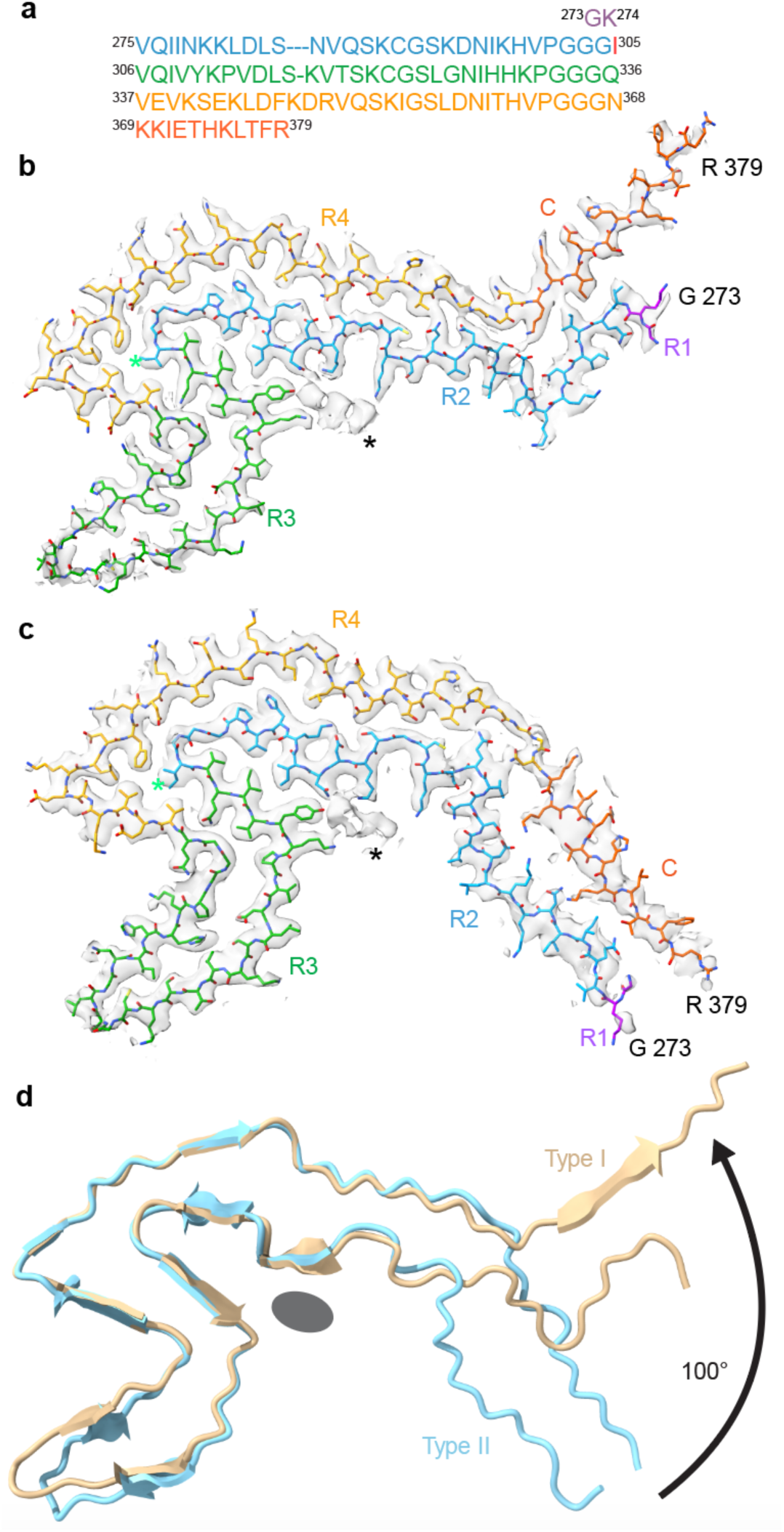
Cryo-EM atomic models and comparison of Type I and Type II S305I folds. **a,** Amino acid sequence and color scheme of the tau structural core that includes R1–R4 residues 273–368 (colored magenta, blue, green, and gold, respectively), and C-terminal residues 369-379 (orange) with S305I highlighted in red. **b-c,** Cryo-EM filament cross-section map/models of Type I (**b**) and Type II (**c**), colored by domain as in (**a**) with unassigned density shown for both filament types (*). **d,** α-carbon backbone overlay of Type I (beige) and Type II (light blue) folds showing 100° rotated position of the R2-C arm and overlapping position of the unassigned density (grey oval).

### Isoleucine Mutation and Additional Density Interaction Support a Structural Basis of the S305I Tau Fold

Based on the related neuropathological features between AGD and *MAPT S305I* FTLD-tau^37^, we next compared the S305I tau structures with the previously published AGD structures^10^. Focusing on the predominant Type I folds, alignment reveals substantial shifts in the overall structure and a more elongated solvent-exposed fold for S305I (**Fig. 4a, Supplementary Video 1**). Beyond these limited similarities, the R3 hairpin, comprising residues 312-334, is the only conserved region between the sporadic AGD and S305I structures. Residues 318-320 and 328-330 form a steric zipper in each fold, and while the relative orientations are substantially rotated, the RMSD is <2.0 Å between the two structures (**Fig. 4a, Extended Data Fig. 2**).

**Figure 4:**
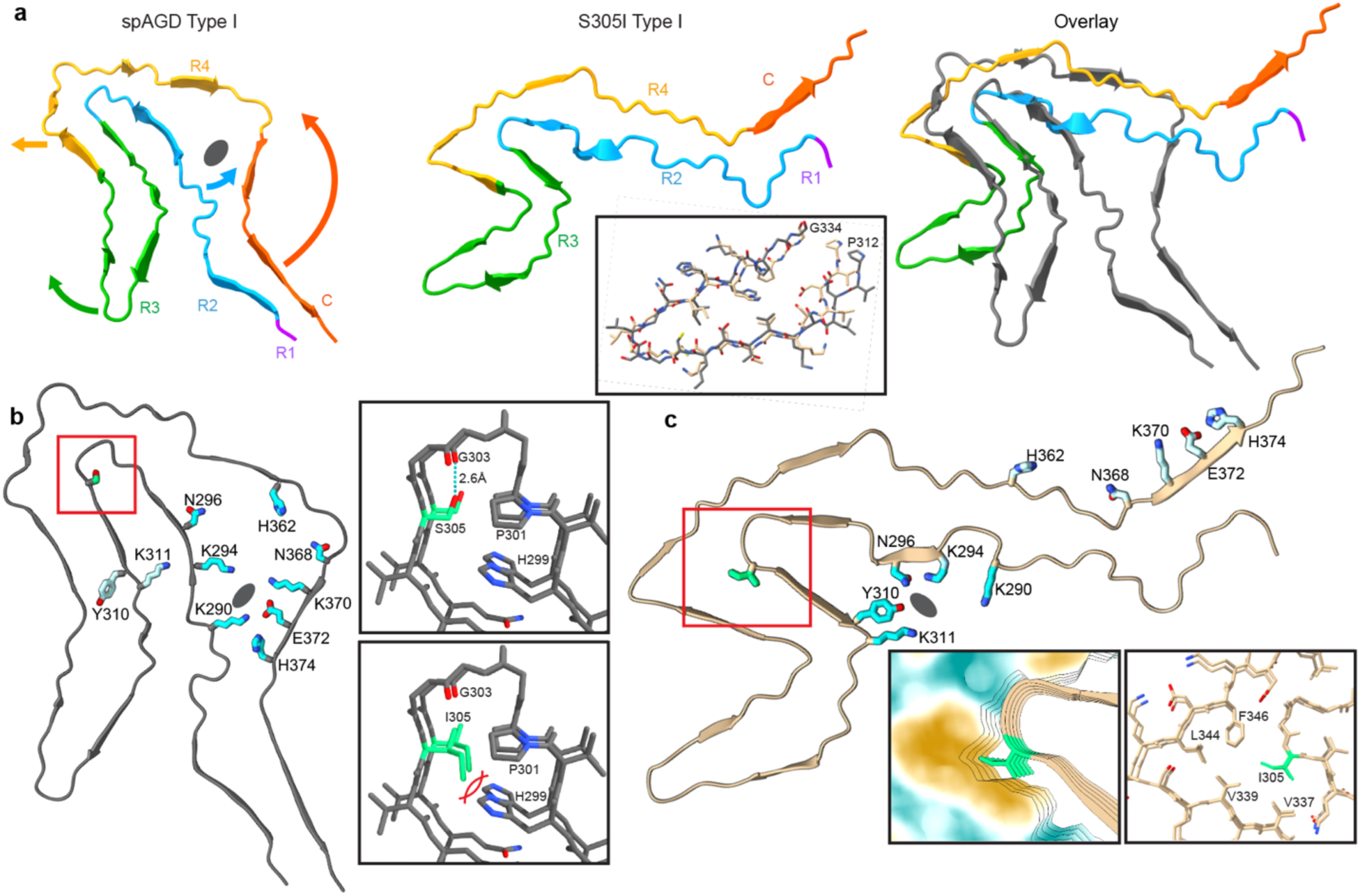
Structure comparison and organization at position 305 for sporadic AGD and S305I Type I filament folds. **a,** The spAGD Type I fold (left), colored as in Fig. 2a with arrows indicating major rearrangements in comparison to S305I Type I (middle), and an overlay of the two structures aligned to R4, with AGD in grey. **b,** The spAGD structure showing solvent-inaccessible polar residues (teal) that form a cavity around an unassigned density (grey oval), and the S305 position (red square). Top inset shows S305 (green) hydrogen bonding with G303; bottom inset shows the incompatibility of a theoretical I305 arrangement in the AGD fold due to a clash with H299. **c,** The S305I structure showing the same polar residues are rotated outward and solvent-accessible, with K294, N296, Y310 and K311 interacting with the unassigned density (grey oval) bound to the outside of the filament. The I305 region is identified (red square) and enlarged with a surface view showing the Ile position in a hydrophobic cleft (left inset) and the surrounding pocket formed by residues V337, V339, L344, and F346 (right inset).

Notably, the large cavity, comprising R4-C and R2 in the AGD fold is collapsed in the S305I structure with no additional internal density in this region (**Fig. 4a**). This enables more continuous side chain interactions between R2 and R4, thereby elongating this stem region. Moreover, in the S305I structure charged residues K290, K295, and N296 in R2 are reoriented toward the solvent-exposed face, where they, together with Y310 and K311, coordinate the additional density located adjacent the R2–R3 junction (**Fig. 4b-c**). These interactions would be incompatible with R2-R3 contacts observed in the AGD structure and thus likely contribute to the rotated position of the R3 hairpin (**Fig. 4a**). Addtionally, several charged residues in the C domain including K370 and E372 that are adjacent to the additional density in the AGD strucure, are rotated and solvent exposed in the S305I structure (**Fig. 4b-c**).

The 305 position lies at the junction of the R2 and R3 domains. In sporadic AGD, S305 is located in a small Gly-rich turn comprised of residues H299-S305 and appears to hydrogen bond with G303 in the adjacent strand (**Fig. 4b, top inset**). Model-building indicated that Ile is physically incompatible with this conformation, eliminating the interaction with G303 and sterically clashing with H299 (**Fig. 4b, bottom inset**). Conversely, in the S305I structure, I305 faces into a hydrophobic groove in the center of the 4-layer fold created by well-packed hydrophobic residues V337, V339, L344, and F346 (**Fig. 4c, insets**). Importantly, these comparisons reveal that Ile mutation likely contributes substantially to the overall fold we observe for the S305I structures.

Considering the large number of tau filament folds that have now been determined and classified based on structural and neuropathological features^10^, we next compared the S305I fold to other disease-associated folds. To this end, we developed a comparison method that evaluates the backbone RMSD between two aligned structures (e.g. S305I Type I and sporadic AGD) over sliding windows of increasing sequence length, thereby enabling the identification of regions with local structural similarity (See Methods). In addition to generating RMSD values for all residue ranges, the results are visualized as triangular heat-maps that highlight regions of structural correspondence across all possible alignments between the two folds (**Extended Data Fig. 3**).

To identify tau filament structures most similar to the S305I Type I fold, we calculated localized RMSD values for all currently available disease-associated tau folds (**Table 2**), where localized RMSD is calculated as the average RMSD value over all residue ranges in the structure. The comparison also uses sequence alignment for all sequentially identical (or similar in the case of mutants) pairs of structures, allowing different splicing forms to be compared. Unexpectedly, the AGD fold showed only moderate similarity, whereas the MAPT P301T Type I^30^ and sporadic CBD folds^13^ exhibited the greatest overall structural resemblance to the S305I fold (**Extended Data Fig. 3**). Analysis of the heatmaps reveals distinct conserved motifs, including the R3 hairpin (residues ∼312-334) and β-strand regions within R2 (∼285-304) and R4-C (∼354-375) (**Extended Data Fig. 3b-c,e**). Among these, the R3 hairpin represents the most consistently conserved structural element, present in AGD Type I, P301T Type I, and CBD folds (**Extended Data Figs. 4**).

Solvent-accessible regions of R2 and R3 in both S305I filament types encase a large well-defined density which extends along the length of the filament axis (**Fig. 5)**. While the individual rungs of the density are well-defined, it contains connected regions parallel to the filament axis. This density is positioned in a positively charged cleft flanked by residues K290, K294, N296, Y310, and K311, indicating it is primarily negatively charged (**Fig. 5a-b**). Given that it must satisfy multiple positive charges in each strand, and from the cross-sectional view does not resemble a polypeptide, we hypothesize that this density is a non-proteinaceous cofactor. The cofactor site (residues 290–311) is highly conserved between Type I and Type II filaments (RMSD = 0.66 Å), which indicates that the co-factor may also be conserved across the two filament types. This conserved region extends from the middle of the R2 region to nearly the end of the R4 region (residues 290–364), with only small differences in backbone bending in the Z-plane (RMSD = 1.97 Å).

**Figure 5:**
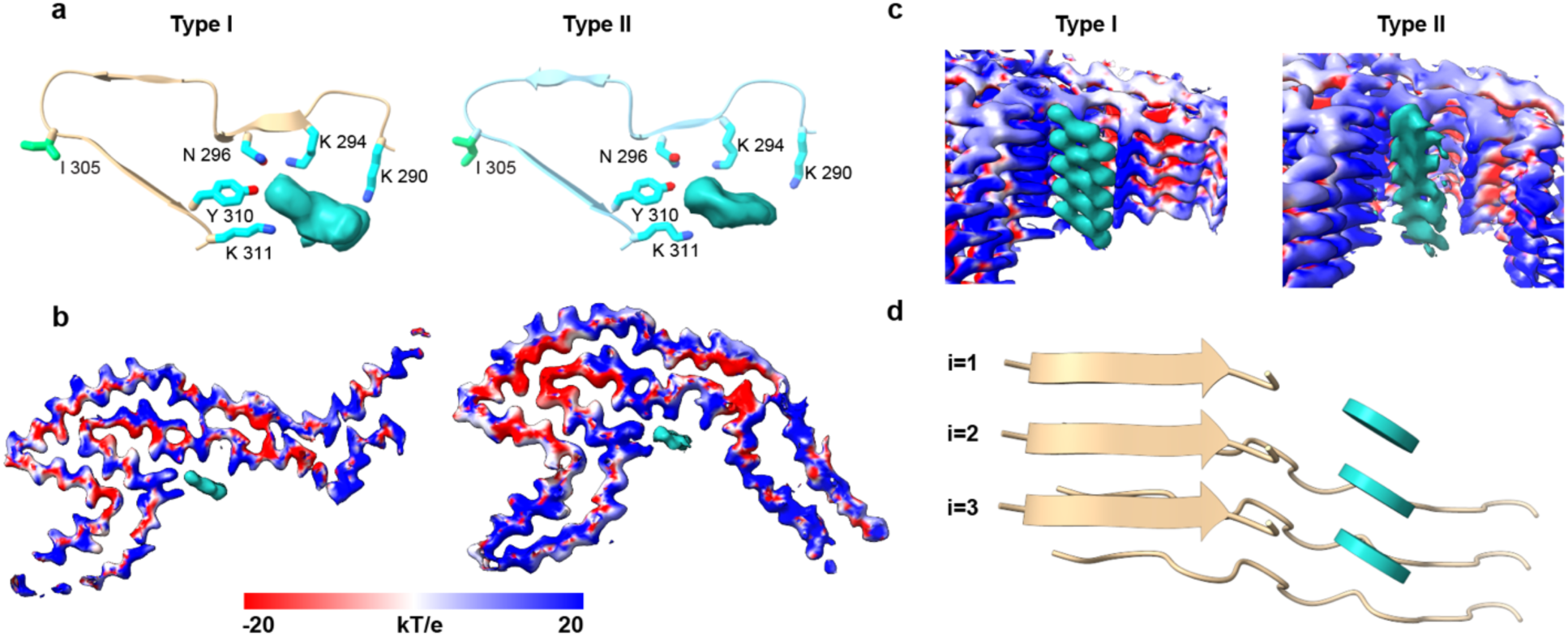
Comparison of unassigned density in both S305I folds; cryo-EM density of S305I Type I (beige) and Type II (light-blue) filaments highlighting the unassigned density along the fibril axis. The unassigned density, which may house a shared cofactor, is shown in cyan. **a,** Top views of the S305I Type I (top) and Type II (bottom) filament residues surrounding the unassigned density. **b,** Top-down views of the unassigned density in the S305I Type I (top) and Type II (bottom) density maps, with electrostatic surfaces colored blue and red to indicate positively and negatively charged regions, respectively. **c,** Side views of the unassigned density in the S305I Type I (top) and Type II (bottom) density maps, with electrostatic surfaces colored blue and red to indicate positively and negatively charged regions, respectively. **d,** Side view of the predominant S305I Type I fibril and a planar representation of the hypothesized cofactor within the unassigned density along the length filament.

Unaccounted, likely non-proteinaceous densities have been observed in several cryo-EM maps of other pathological tau fibrils^10,12,13^. These have been attributed to anions, hydrophobic or charged metabolites or negatively charged RNA or polyphosphate that bind to complementary regions of the fibril, potentially stabilizing the fold^10,12,13,44–46^. Thus, there is growing interest in identifying these molecules as contributors to distinct tau folds, however the densities are often diffuse and low-resolution, making their interaction difficult to define. Notably, based on our RMSD analysis above, we identify the that P301T Type I tau fold contains a region of strong structural homology compared to the S305I fold that encompasses the R2–R3 junction (residues ∼280-314) containing residue 305 and residues K290, K294, N296, Y310 and K311 that coordinate the additional density observed in our maps (**Extended Data Fig. 3e**)^30^. Upon examination of the deposited P301T Type I map (EMD-51320), we identify a strikingly similar well-defined unaccounted density in this region that is coordinated by the same residues we observe in the S305I structures (**Extended Data Fig. 5**). We note this density was not previously reported for the P301T Type I fold^30^. Based on these similarities, we propose that the additional density in both structures likely corresponds to the same bound cellular metabolite or polyanion. Together, these findings reveal structural commonalities between S305I and other disease-associated tau folds and suggest that both the S305I substitution and the associated bound factor together contribute to the distinct structural fold observed in S305I.

### Molecular modeling and In vitro analysis indicate polyanions and S305I mutation contribute to tau fibrilization

Based on the well-defined stacked arrangement of the extra density and positive charge of the coordinating residues in the S305I structure, we next docked several candidate molecules into the density (see Methods), to better understand potential cofactor interactions (**Extended Data Fig. 6**). Based on the size of the density and surrounding residues, we docked polyphosphate in addition to RNA and inositol diphosphate^47^, a common cellular metabolite containing two anionic phosphate groups, into the density. Inositol diphosphate fits well in an arrangement in which the phosphate groups interact with K290, K294, Y310, and K311, while the inositol rings stack together along the fibril axis with a geometry that matches the overall 4.8 Å repeat of the amyloid structure. A polyphosphate chain can dock into the density in straight or bent configurations with the phosphate groups interacting with K294 and K311, but the fit is incomplete with the chain accounting for only a fraction of the available density (**Extended Data Fig. 6b-c**). We next docked poly-A and poly-G strands as a model for RNA. Poly-G fits well in a single arrangement in which the phosphodiester backbone packs along the fibril groove, interacting with K294, Y310, and K311, while the purine ring projects outward at an angle that matches the density and enables an interaction with K290 (**Extended Data Fig. 6d-g**). This stacked ring arrangement is similar to previous amyloid-small molecule co-structures, in which pi-pi stacking between the aromatic rings of each ligand supports interactions with the fibril^31,32,48^. We note that in all modeling tests, the negatively-charged phosphodiester linkage is positioned in the cationic groove of the S305I fibril, where it forms complementary electrostatic interactions with adjacent Lys residues. Thus, this analysis further supports a direct role for polyanions such as RNA or phosphate-containing metabolites functioning as cofactors that stably bind pathogenic folds of tau.

Considering the Ile residue appears to directly organize the 4-layer tau fold of the S305I structures we determined (**Fig. 4**), we hypothesized that the mutation itself may contribute to disease pathology by promoting tau aggregation in addition to its reported splicing defects and increased 4R tau expression^36,37^. To assess the impact of S305I on tau aggregation, we compared the *in vitro* fibrilization kinetics of purified, recombinant 0N4R tau containing S305I or P301S, a well-characterized familial mutation known to enhance aggregation, to wildtype, normal tau using an established Thioflavin T (ThT) fluorescence assay^49,50^. Here we utilized polyphosphate, at 0.125 mg/mL, as an established polyanion inducer of fibrilization^51^ and determined the overall fibrillization rate and aggregation propensity (as the inverse fluorescence half-time (t_1/2_))^52^, as described. (**Fig. 6a and Extended Data Fig. 7**). Under these conditions, we observe that the S305I fibrilization rate is significantly faster (by nearly 2-fold) compared to wildtype and slightly faster than P301S (**Fig. 6b)**. Additionally, based on the aggregation propensity calculation, S305I is modestly faster than wildtype while P301S is substantially faster (nearly 3-fold) **(Fig. 6c)**. From this analysis, we conclude that the presence of the S305I mutation indeed promotes tau aggregation, similar to what has been observed for other aggregation-prone mutations such as P301S. Finally, we assessed the polyphosphate-induced S305I tau fibrils by cryo-EM and identify they form intact twisted fibrillar structures that appear distinct from other *in vitro* fibrils^53^ (**Fig. 6d)**. Upon 2D classification analysis we identify these fibrils are heterogeneous and adopt several different folds which prevented 3D structure determination. However, several classes show a distinct curvature that appears similar to the wavy morphology we observe for the brain-derived S305I fibrils indicating some structural similarities may be present between these folds (**Extended Data Fig. 7d)**.

**Figure 6:**
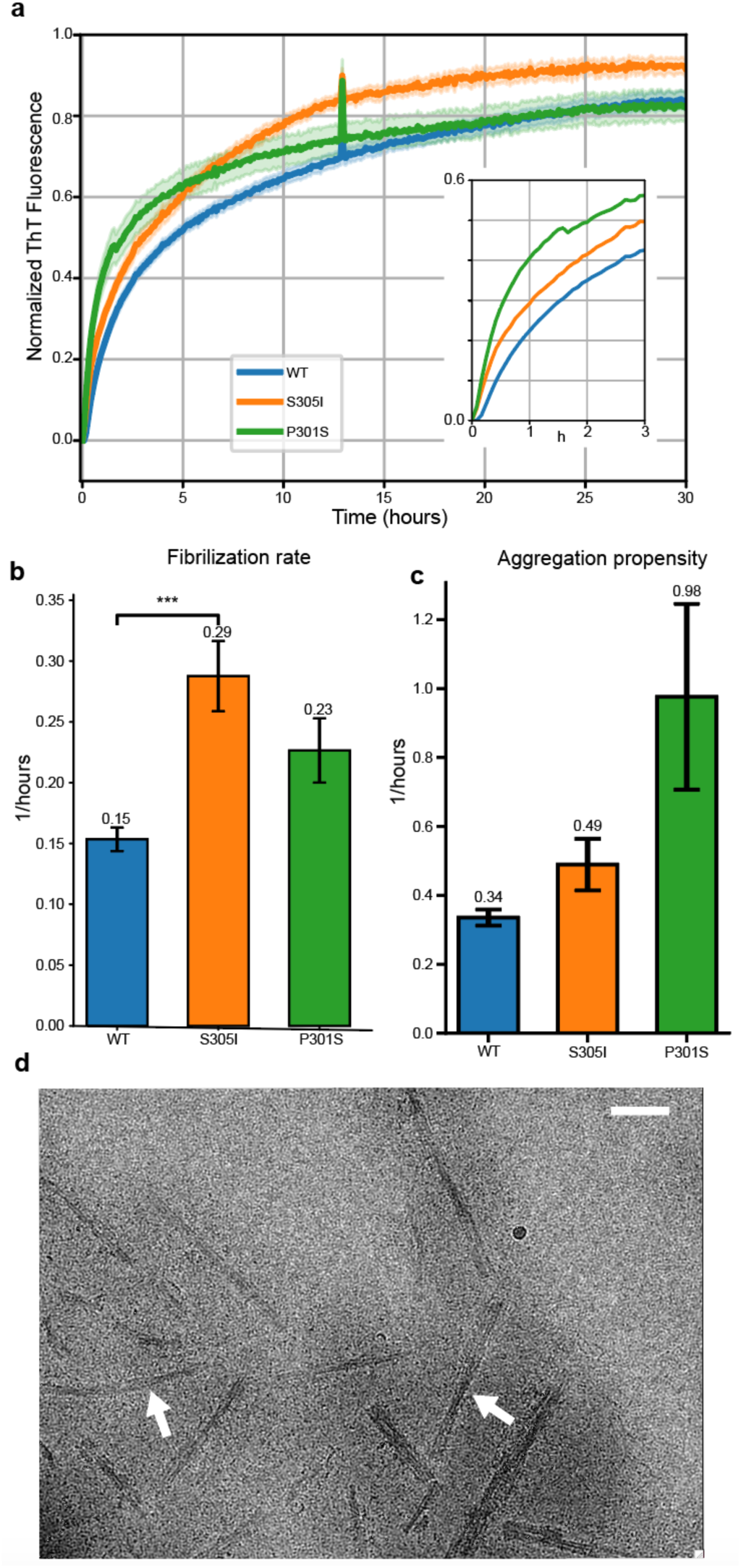
Increased *in vitro* fibrilization of S305I tau. **a,** Fibrilization curves for WT (blue), S305I (orange) and P301S (blue) 0N4R tau, using polyphosphate as the inducer. **b,** Filament elongation rate, calculated from the Gompertz function (see Methods). S305I is significantly faster than WT, as determined by a one-way ANOVA with Dunnett’s post-hoc test. **c,** Aggregation propensity for each construct, calculated as the inverse t_1/2_. **d,** Representative cryo-electron micrograph of recombinant S305I tau following polyphosphate-induced aggregation showing well-resolved twisted filaments (arrows) (Scale bar = 50 nm).

## Discussion

Pathogenic mutations in the *MAPT* gene continue to be identified, with currently over 60 linked to FTLD^54^. Many give rise to neuropathological phenotypes that closely resemble those observed in well-characterized sporadic tauopathies, including AD, PSP, CBD, and others^55^. On this basis, it has been proposed that distinct *MAPT* mutations can be classified according to their corresponding sporadic tauopathy counterparts^24^. However, familial tauopathies often show earlier onset and greater severity than sporadic forms, potentially driven by enhanced tau aggregation, excess 4R tau expression due to intronic or silent mutations that disrupt intron 10 splicing, or other mechanisms^56–60^. Consequently, the influence of specific tau mutations on tau structure, aggregation behavior, and disease phenotype remains unclear. Here we determined tau filament structures from postmortem tissue of a patient with a rare *MAPT* familial FTLD mutation, S305I, which resides at the exon 10 splice junction and is associated with severe early-onset FTLD with pathological features that resemble AGD. Our structures reveal tau S305I filaments adopt two related, but previously unreported 4-layer paired filament folds involving tau repeat domain residues 273-379. The structures are distinct from sporadic AGD, which is notable because of the related neuropathology including the presence of tau deposits in the form of argyrophilic grains (**Fig. 1**). The Ile sidechain in the S305I fold is efficiently packed in a hydrophobic cleft; the more polar Ser sidechain in the sporadic AGD fold would be less well packed nor would it be hydrophobically stabilized. The structure also contains an unidentified but well-defined density that stacks along an outer, positively charged cleft, potentially also contributing to the unique fold. Notably, molecular modeling indicates RNA or other phosphate-containing small-molecules could bind favorably to the outer cleft and potentially account for the unidentified density, as has previously been proposed for other tau structures^10,44,45^. Importantly, these structures add to the diversity of tau folds that form in disease and reveal how a single mutation may contribute mechanistically to FTLD.

Disease mutations at the 305 position and other sites in intron 10 are reported to disrupt the RNA stem-loop structure at the splice junction, resulting in reduced splicing and an overproduction of 4R tau relative to 3R^34,61^. However, differences in the clinical and pathological features occur with these mutations, indicating that overproduction of 4R tau alone is not a predictor of specific pathology^61^. The *MAPT S305I* mutation, like other 4R tau–associated tauopathies, presents with early-onset symptoms and rapid progression^21,37,62^ but exhibits AGD pathology distinct from other substitutions. S305N is linked to Pick-like bodies composed of 4R tau^63^, while the synonymous S305S (AGT→AGC) shows PSP-like pathology^64^. Thus, these differences further indicate that disease variation arises from mutation-specific effects beyond 4R tau overproduction. Moreover, based on our structures, we predict S305N is unlikely to form the S305I tau fold (**Extended Data Fig. 8**) due to asparagine’s polar side chain pointed inward toward the apolar pocket where I305 resides. Additionally, the S305N mutation is also incompatible with the sporadic AGD fold, due to a 2.5 Å steric clash the asparagine’s most favorable rotamer makes with the H299 imidazole ring in chain *i+1* directly below. Thus, we predict S305N likely adopts an additional unknown fold, that is associated with distinct pathology and disease characteristics^30^.

Structural determination of patient-derived tau filaments for 5 other familial *MAPT* mutations has been achieved thus far. Some adopt folds previously identified in sporadic tauopathies (ΔK281, V337M, R406W)^14,22^, while others have unique folds (P301L and P301T)^30^. Of the 3 familial disease folds that match sporadic diseases, only V337M is located in the ordered filament core, and this mutation is both sterically and electrostatically tolerated by the AD PHF fold^14^. In the P301L and P301T folds, which do not match those of any sporadic diseases, the mutation of proline allows for an additional hydrogen bond, potentially driving the formation of an alternative fold. The central position of S305 in the 4-layer AGD fold and the destabilizing steric clash the mutant I305 has with H299 in R2 (**Fig. 4b**) suggests that this mutation may play a more substantial role in driving a specific tau fold compared to other mutations. To further understand the relationship between tau mutations and fibrilization, we induced aggregation of WT, S305I, and P301S tau via the addition of polyphosphate. S305I has a higher propensity to aggregate and fibrilization rate relative to WT, further evidence the mutation itself may act as a distinct driver of aggregation and the structural fold^14,22,30^. How the specific tau fold and increased aggregation propensity of S305I mechanistically contribute to the neuropathology and early disease onset characteristics of this FTLD remains to be determined.

As there is often clinical and neuropathological overlap between familial and sporadic forms of FTLD, the question as to whether these are familial versions of sporadic tauopathies remains open and has important implications for the design of models, diagnostics, and therapeutics for FTLD. Neuropathologically, S305I results in argyrophilic grains resembling those found in AGD, which led to the hypothesis that this FTLD might be a familial form of AGD^37^. Notable differences between FTLD-tau/*MAPT* S305I and sporadic AGD, however, raised the possibility that they represent distinct diseases. The pathology of sporadic AGD is primarily found in the limbic system, whereas S305I tau inclusions extend much more widely. Glial characteristics also differ, with sporadic AGD showing more prevalent granular-fuzzy type astroglial cytoplasmic inclusions. Recently, comparison of filament folds has been suggested as a metric for classification of tauopathies^10^. Given this, our structural findings demonstrate that S305I FTLD-tau is distinct from AGD, as opposed to an early-onset form of the disease. As no broad structural relationship has been established between FTLD-tau/*MAPT* and sporadic tauopathies, further investigation is warranted regarding the classification of these diseases.

Many amyloid filament structures from patients with tauopathies have shown additional non-proteinaceous densities, indicating that the specific tau fold and its stability may require the binding of distinct small-molecule metabolites or polyanions^10^. In Type I and II sporadic AGD folds, a small, poorly resolved density is present in a solvent-protected groove, interacting primarily with positively charged residues K294 and K370. In S305I, both folds exhibit a collapse of this groove and the R2 strand with these residues is flipped, introducing new sidechain interactions with R3 (**Fig. 4b-c)**. Notably, this R2-R3 interaction permits the formation of the solvent-exposed cleft in which K290, K294 and N296 in R2, and Y310 and K311 in R3 together contact the exogenous molecules seen as stacked density in the S305I folds. Interestingly, these same 5 residues in the P301T fold form a very similar cleft that houses unassigned density which strongly resembles that in the S305I folds^30^. Unassigned densities in these and other tauopathy filament folds raise questions about the role and timing of these densities with respect to filament formation. Upstream effects on cellular homeostasis could be driving specific filament folds; alternatively, filaments could be sequestering small molecules after amyloid formation. Irrespective of the temporal sequence, binding of a highly anionic molecule would be expected to thermodynamically stabilize the conformation found in S305I fibrils. Previous work has speculated that some of the unassigned densities in tau and other fibril structures are associated with anionic molecules^10,65–68^, including RNA^44^. This is consistent with our modeling, which shows that multiple RNA molecules and other anions fit the unassigned density in the S305I folds. Thus, it is of interest to identify the anionic cofactors and elucidate their role in fibril formation and stabilization.

## Materials and Methods

### Ethical review process and informed consent

Human brain tissue samples were obtained from the Neurodegenerative Disease Brain Bank at the University of California, San Francisco. Prior to autopsy, participants or their surrogates provided informed consent for genetic testing and for brain donation, in accordance with the principles of the Declaration of Helsinki. Procedures were approved by the UCSF Committee on Human Research.

### Case materials

Genetic screening was performed for common and rare FTD-related genes as previously described^40^. Patient samples were screened by targeted sequencing of the seven most common dementia-associated genes (microtubule-associated protein tau [*MAPT*], progranulin [*GRN*], TAR DNA binding protein [*TARDBP*], fused in sarcoma [*FUS*], amyloid beta precursor protein, presenilin 1 [*PSEN1*], and presenilin 2 [*PSEN2*], as described previously^40^. Coding and exon-intron boundary regions of these seven genes were screened for known or novel (likely) pathogenic variants. Variants identified in the screening were confirmed by Sanger sequencing^40^.

Neuropathological diagnoses were made following consensus diagnostic criteria^69–71^ using previously described histological and immunohistochemical methods^72–74^. Briefly, eight-micrometer thick sections were cut from formalin-fixed paraffin-embedded blocks and stained with hematoxylin & eosin and with antibodies against hyperphosphorylated tau (CP13, anti-mouse, 1:250, targeting pSer202, gift from Peter Davies, Feinstein Institute for Medical Research, USA), amyloid-beta (anti-mouse, 1:500, Millipore, MAB5206), TDP-43 (anti-rabbit, 1:4K, ProteinTech, 10782-2), alpha-synuclein (anti-mouse, 1:5K, LB509, gift from John Trojanowski and Virginia Lee, University of Pennsylvania, USA), and 3R-tau (anti-mouse, 1:2K, Millipore, 05-803) and 4R-tau (anti-mouse, 1:2K, Millipore, 05-804), as described previously^72^. Cases were identified based on neuropathological diagnosis and genetic screening. Photomicrographs of the hyperphosphorylated tau-stained slides from the inferior frontal gyrus and inferior temporal gyrus were captured using a digital camera (Nikon Digital Sight DS-Fi1) mounted on a Nikon Eclipse 80i microscope using NIS-Elements (Version 3.22, Nikon) imaging software. Fresh-frozen blocks of the inferior frontal gyrus and inferior temporal gyrus were cut from freshly frozen coronal slabs that had been stored at -80 °C.

### Experimental design

Samples were selected based on brain tissue availability and post-mortem neuropathological examination. Sex was not considered in this study, as the structure of AGD filaments does not vary by sex. Randomization and blinding were not performed in this study, as there was only one patient available at the NDBB carrying the *S305I MAPT* mutation.

### Tau Filament extraction

Sarkosyl-insoluble material was extracted from the inferior frontal cortex using a protocol similar to previous amyloid filament purifications^10^. In brief, tissues were homogenized in 20 volumes (v/w) of extraction buffer consisting of 10 mM Tris-HCl, pH 7.5, 0.8 M NaCl, 10% sucrose and 1 mM EGTA. Homogenates were brought to 2% sarkosyl and incubated for 30 min at 37 °C. After a 10-min centrifugation at 20,000g at 4 °C, the supernatants were spun at 100,000g at 4 °C for 20 min. The pellets were resuspended in 700 μL/g extraction buffer and centrifuged at 9,500g for 10 min. The supernatants were diluted threefold in 50 mM Tris-HCl, pH 7.5, containing 0.15 M NaCl, 10% sucrose and 0.2% sarkosyl, and spun at 166,000g for 30 min. Sarkosyl-insoluble pellets were resuspended in 25 μL/g of 20 mM Tris-HCl, pH 7.4 and 100 mM NaCl and used for cryo-EM.

### Negative stain imaging

Purified frontal cortex tissue was diluted 1:10 for a final concentration of 150 μL/g frozen tissue. 3 μL was added to a glow-discharged 400 mesh copper grid with a layer of amorphous carbon. After 30 seconds, the grid was blotted with filter paper, washed, and blotted twice with water. 4 μL of 0.75% uranyl formate was then added and blotted. Three more 4 μL aliquots of uranyl formate were added and removed by vacuum aspiration. Images were collected on a Talos L120C (Thermo Fisher Scientific) operating at 120 kV and equipped with a Ceta-D (Thermo Fisher Scientific) camera.

### Cryo-EM grid preparation and data collection

Extracted tau filaments were incubated at 37 °C in a 0.4 g/mL pronase solution for 30 minutes and centrifuged at 3,000 g for 5 minutes, then applied to glow-discharged holey carbon grids (Quantifoil Au R1.2/1.3, 200 mesh). Then, 3 μL of purified tau filaments were plunge-frozen in liquid ethane using a Vitrobot Mark IV System (Thermo Fisher Scientific). Images were acquired on a 300 KeV Titan Krios microscope (Thermo Fisher Scientific) equipped with a K3 direct electron-detection camera (Gatan, Inc.) with a BioQuantum energy filter (Gatan, Inc.) with a slit width of 20 eV. Super-resolution movies were recorded at 105,000x magnification (pixel size: 0.417 Å/pixel) with a defocus range of 0.8 to 1.8 µm and a total exposure time of 2.024 seconds fractionated into 0.025-second subframes.

### Image Processing

Datasets were processed in RELION using standard helical reconstruction^75^. The movies were motion-corrected using MotionCor2^76^ and Fourier-cropped by two to give the final pixel size of 0.834 Å per pixel. The dose-weighted summed micrographs were directly imported to RELION^77,78^ and used for further processing. Contrast transfer function (CTF) was estimated using CTFFIND 4.1^79^. The filaments were picked manually and extracted with a box size of 900 pixels downsampled to 300 pixels. 2D classification was used to separate filamentous segments from junk particles, and then additional 2D classification was performed to distinguish Type I from Type II filament. 3D initial models were then created *de novo* from the 2D class averages using the crossover distances measured from the 2D class averages, 600 Å for type I and 750 Å for type II, using RELION’s *relion_helix_inimodel2d* feature^80^. Particles associated with each filament type were re-extracted separately using a 300 pixel box, and 3D classification was performed using each initial model low-pass filtered to 15 Å. The best classes were selected from each 3D classification, and the final rise and twist was optimized in 3D auto-refinement. The final 3D densities were sharpened using standard postprocessing methods in RELION, and the final resolution was determined from Fourier shell correlation at 0.143 Å from two independently refined half-maps using a soft-edged solvent mask. The final resolution of the Type I filament was 3.1 Å and the resolution of the Type II filament was 3.2 Å.

### Model Building and Refinement

The atomic model of the S305I and normal AGD structures were docked manually into the density using ChimeraX^81^ and ISOLDE^42^, using predicted models from ModelAngelo^43^ as a guide. The model was refined against the density using Phenix. Figures were prepared using ChimeraX, PyMOL, and ABPS^82^.

### Filament Fold Comparison

Code was generated that evaluates the backbone RMSD between two filament folds over sliding windows of increasing sequence length. Fully annotated code is available at https://github.com/degrado-lab/amyloid_analysis.

### Docking of Polyanions

Docking studies were performed using the Molecular Operating Environment (MOE; Chemical Computing Group, Build MOE 2024.0601). The protein structure was derived from the S305I Type I fold. The poly(A) RNA ligand was obtained from PDB entry 8ZWJ. A poly(G) RNA model was generated by modifying the purine base of the poly(A) RNA structure. Linear polyphosphate models were constructed in MOE, and a helical polyphosphate model was generated by manually building the RNA phosphate backbone followed by energy minimization. The inositol diphosphate structure was obtained from PDB entry 1I9Z^83^. Docking was carried out using the MOE protein-protein docking module, in which the ligand structures were sampled against the protein target. Candidate complexes were generated through rigid-body placement followed by refinement and scoring. The resulting docking poses were clustered base on interface interaction fingerprints, and representative structures from the top-ranked clusters were selected for further analysis.

### Recombinant protein expression and purification

Recombinant 0N4R tau constructs were expressed from a pET-28a vector and transfected into *Escherichia coli* BL21(DE3) plysS competent cells. Starter cultures containing a single colony of transformed cells were grown in Terrific Broth (TB) containing 50 μg/mL kanamycin and 25 μg/mL chloramphenicol overnight (16-18 hours) at 37°C with constant shaking at 200 rpm. Then, 15 mL of the starter culture was used to inoculate 1 L of terrific broth (TB), containing 50 μg/mL kanamycin and 25 μg/mL chloramphenicol. Cell cultures (4 L per construct) were grown at 37°C with constant shaking at 200 rpm until an optical density (OD_600_) of 1.0 was reached. Expression was induced with 1 mM IPTG for 16 hours at 16°C, then harvested by centrifugation (4000 rpm for 20 minutes) and the cell pellets were flash frozen in liquid nitrogen.

To purify tau, cell pellets were resuspended in ice-cold lysis buffer (50mM MES pH 6.0, 20 mM NaCl, 2 mM DTT), supplemented with two cOmplete EDTA-free protease cocktail inhibitor tablets (Roche). Cells were lysed by dounce homogenization and sonication, and then boiled for 20 minutes at 80°C. The boiled lysate was clarified via centrifugation at 30,000 g for 30 minutes. The clarified lysate was filtered using a 0.2-0.3 micron syringe filter and loaded onto a Cytiva CaptoS 5 mL column (GE healthcare) equilibrated in a low salt buffer (50 mM MES pH 6.0, 20 mM NaCl, 2 mM DTT). The sample was eluted using a 16-column volume (CV) gradient from 0% to 100% high salt buffer (20 mM MES pH 6.0, 1.5 M NaCl, 2 mM DTT). Protein-containing fractions were pooled and concentrated before loading into a 5 mL loop and injecting into a HiLoad Superdex 200 16/600 column equilibrated with 1x PBS pH 7.4, 5% glycerol, 2 mM TCEP for size-exclusion chromatography. Fractions containing protein were concentrated, flash frozen, and stored at -80°C. Purity was analyzed via SDS-PAGE and final concentration was verified via BCA assay (Pierce).

### Tau Fibrillization

Aggregation reactions were performed in 384-well plates (Corning 4511). In each well, purified 10 μM 0N4R tau constructs (WT, P301S, S305I) were added to freshly prepared assay buffer (1x PBS pH 7.4, 2 mM TCEP). Aggregation was induced immediately prior to the start of the assay by adding polyphosphate at a final concentration of 0.125 mg/mL, with each well containing a total volume of 20 μL. Aggregation reactions were performed at 37°C with continuous shaking. Fluorescence measurements (excitation = 444 nm, emission = 475 nm, cutoff = 490 nm) were recorded in a Spectramax M5 microplate reader (Molecular Devices). Readings were taken every 5 minutes for at least 24 hours. Each experiment was performed with 9 replicate wells per condition, with 3 separate replicate experiments run.

### Fibrilization Data Analysis

The first five data points were averaged and this background was subtracted from each sample. Baseline subtracted curves were fit using the Levenberg-Marquardt nonlinear least-squares method to the Gompertz function:

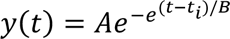

Where y is the fluorescent value at time (t), A is the amplitude of the curve, t_i_ is the inflection point, b is 1/k_app_ where k_app_ is the elongation constant. Individual experimental curves with max raw RFU values less than 50 and a mean-squared error (MSE) higher than 1 standard deviation of the mean of the maximum ThT signal were discarded. Analysis of the discarded curves shows features consistent with bubbles in reaction wells. Final data were plotted in GraphPad PRISM and with Matplotlib.

## Supporting information

AGD to S305I Type I Morph

AGD to S305I Type II Morph

## Data Availability

Cryo-EM maps and atomic coordinates have been deposited in the EMDB and PDB with accession codes: EMD-75195 (Type I) and EMD-75196 (Type II), and PDB 10IJ (Type I) and 10IK (Type II). Filament fold comparison code is available at https://github.com/degrado-lab/amyloid_analysis. Any other relevant data are available from the corresponding authors upon request.

## Acknowledgements

This work was funded by the Rainwater Charitable Foundation (D.R.S.), and the National Institutes of Health (P01AG002132: D.R.S. and W.F.D). The UCSF Neurodegenerative Disease Brain Bank receives funding support from NIH grants P30AG062422, P01AG019724, U01AG057195, and U19AG063911, as well as the Rainwater Charitable Foundation and the Bluefield Project to Cure FTD. Data collection and dissemination of the genetic data presented in this manuscript was supported by the ALLFTD Consortium (U19: AG063911, funded by the National Institute on Aging and the National Institute of Neurological Diseases and Stroke) and the former ARTFL & LEFFTDS Consortia (ARTFL: U54 NS092089, funded by the National Institute of Neurological Diseases and Stroke and National Center for Advancing Translational Sciences; LEFFTDS: U01 AG045390, funded by the National Institute on Aging and the National Institute of Neurological Diseases and Stroke). Additional funding support was provided by the National Institutes of Health (NIH) National Institute on Aging (NIA) for Frontotemporal Dementia: Genes, Images, and Emotions: P01 AG019724 and UCSF Alzheimer’s Disease Research Center: P30 AG062422. Samples from the National Centralized Repository for Alzheimer’s Disease and Related Dementias (NCRAD), which receives government support under a cooperative agreement grant (U24 AG21886) awarded by the National Institute on Aging (NIA), were used in this study. The authors acknowledge the invaluable contributions of the study participants and families as well as the assistance of the support staff at each of the participating sites. H. S. P. received postdoctoral fellowship funding from the BrightFocus Foundation (A141860). We thank Dr. Carlo Condello for his insight into the fluorescence kinetic assay and unassigned density analysis.

## Author Contributions

W.W.S., D.R.S., L.T.G., H.J.R., and M.L.G., and G.E.M conceptualized the study. H.S.P., G.E.M., W.W.S., D.R.S., and W.F.D. designed the experiments. H.S.P. and G.E.M., A.A.M., and E.T. performed cryo-EM sample preparation, data collection, and data processing. H.S.P. generated atomic models on the basis of cryo-EM maps. A.N.L., S.S., E.M.R., A.L.L., and W.W.S. performed neurology, pathology, genetics and histology studies. M.L., H.S.P., A.Y. and A.Q. cloned and purified recombinant tau constructs and performed fibrilization assays. H.J. and H.S.P. performed docking studies. R.C.K. and H.S.P. developed the code for structural homology analysis. H.S.P., G.E.M., A.N.L., W.W.S, and D.R.S. wrote and edited the manuscript.

## Ethics Declarations

W.F.D. is a member of the scientific advisory boards of Alzheon Inc., Pliant, Longevity, CyteGen, Amai, and ADRx Inc., none of which contributed support for this study. W.W.S has received consulting fees from BridgeBio, Guidepoint Global, Inc., and GLG Council; speaker honoraria from Verge Genomics; and compensation as a member of the scientific advisory board for Lyterian Therapeutics, none of which contributed support for this study. The remaining authors declare no competing interests.

