## Extended Data for "Distinct tau filament folds in familial frontotemporal dementia due to the *MAPT* S305I mutation"

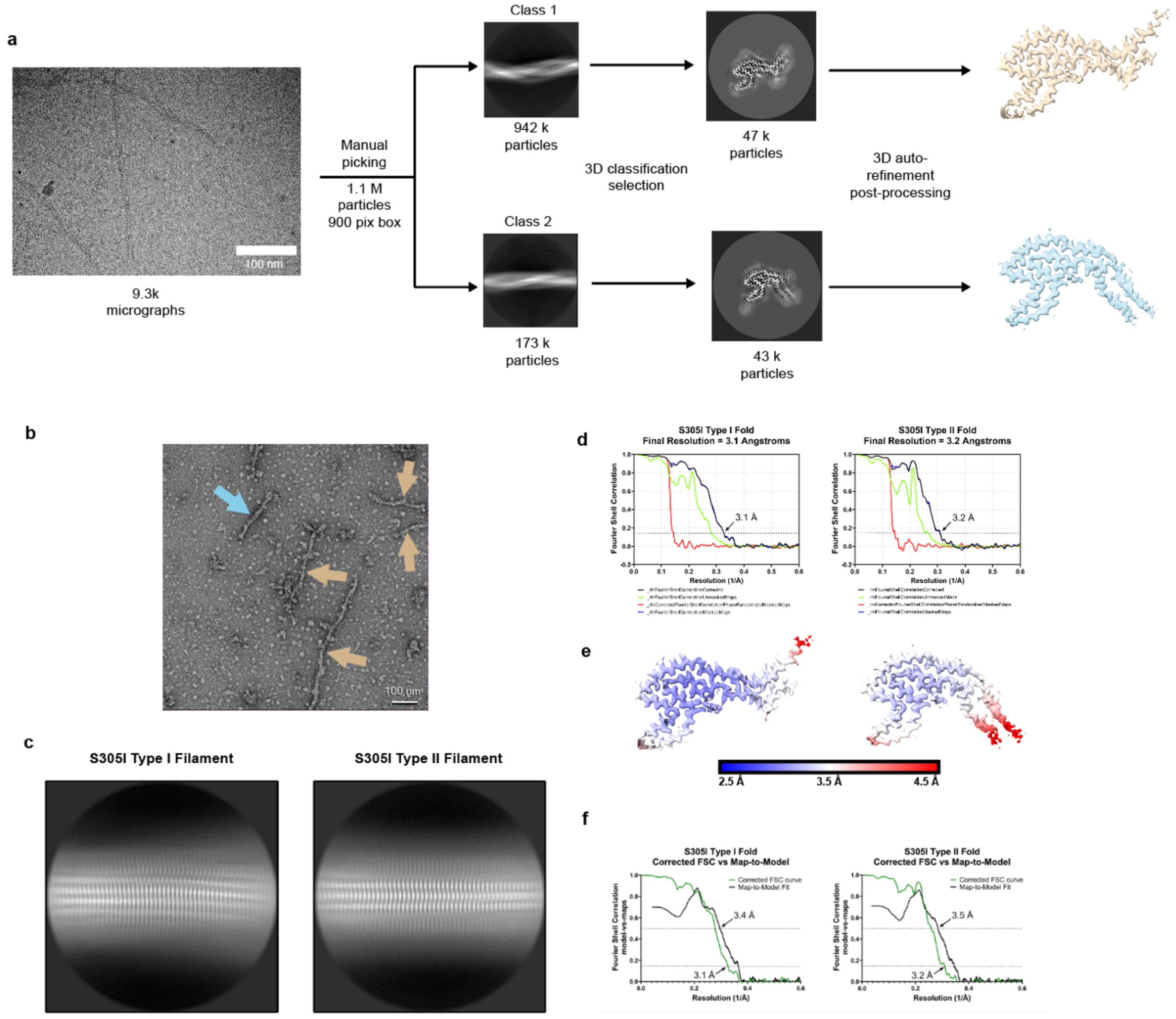

**Extended Data Fig. 1. Cryo-EM overview**. (**a**) Processing workflow for Type I and Type II S305I filaments. (**b**), nsEM of S305I tau filaments purified from patient brain tissue. (**c**), Representative 2D class averages of S305I Type I and II filaments (250 Å box size). (**d**), Fourier shell correlation (FSC) plots for two independently refined cryo-EM half maps of S305I Type I (left) and Type II (right) folds. (**e**), Local resolution map of S305I Type I (left) and Type II (right) folds, showing higher resolution near the core and unassigned density, and lower resolution in the exposed hairpin and tail regions. (**f**), FSC curves for map-to-model fit.

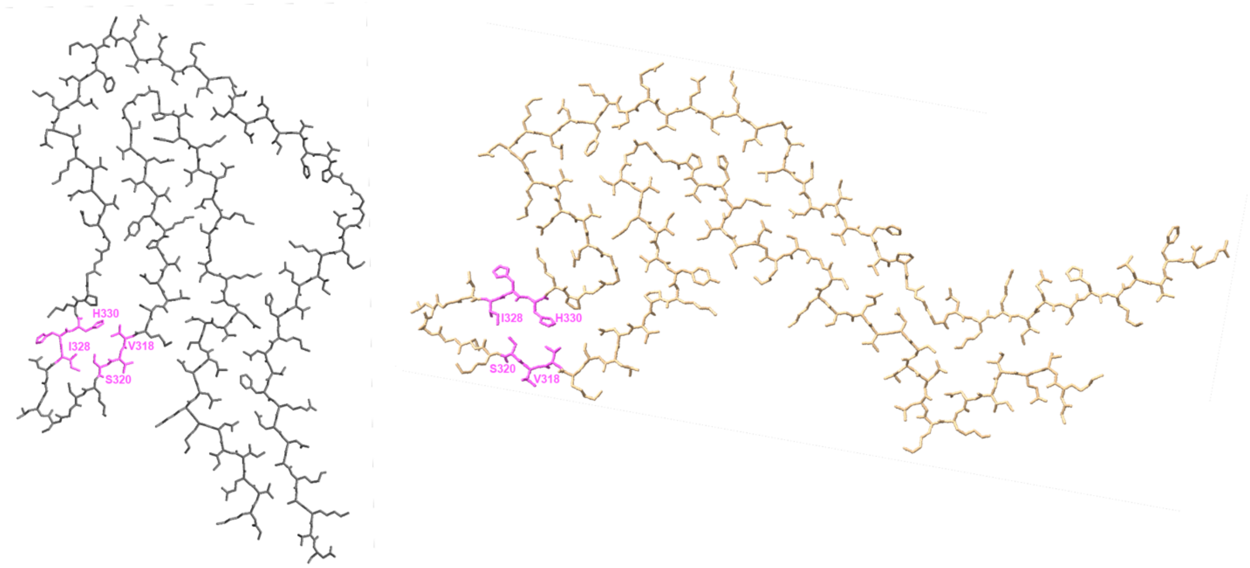

**Extended Data Fig. 2. Sporadic AGD and S305I tau folds share cross-β interactions.** Conserved zipper motif in the R3 hairpin of sporadic AGD (grey; PDB: 7P6D) and S305I (tan) folds.

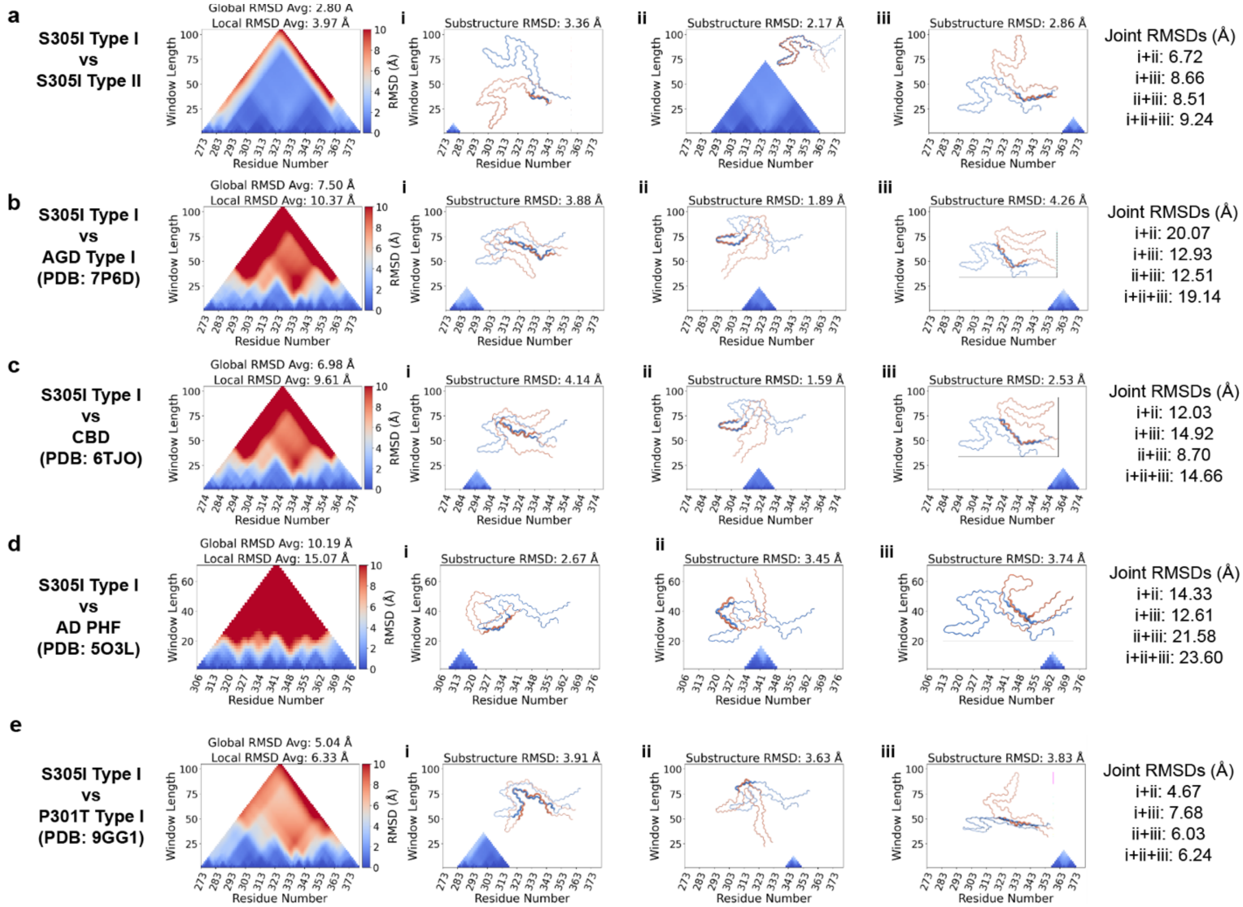

**Extended Data Fig. 3. RMSD comparisons across tau filament folds.** Triangular heat maps measure similarity between filament structures at all ranges, by calculating backbone RMSDs of two tau structures over windows of increasing length, from several-residue segments (bottom row, essentially RMSD = 0 Å, dark blue) to full structures (high RMSD, dark red). The color scale at the right of the triangles shows the RMSD up from 0 to 10 Å, with RMSDs above 10 Å remaining the same dark red for readability. The joint RMSDs at right are RMSD values calculated by traditional selection and global alignment, demonstrating significantly higher RMSD values compared to the locally aligned substructure RMSD values (**a**), S305I Type I fold compared to S305I Type II. (**b**), S305I Type I fold compared to AGD Type I (PDB: 7P6D). (**c**), S305I Type I fold compared to Corticobasal Degeneration (CBD; PDB: 6TJO). (**d**), S305I Type I fold compared to Alzheimer's disease paired helical filament (PHF) (AD; PDB: 5O3L). (**e**), S305I Type I fold compared to P301T Type I (PDB: 9GG1).

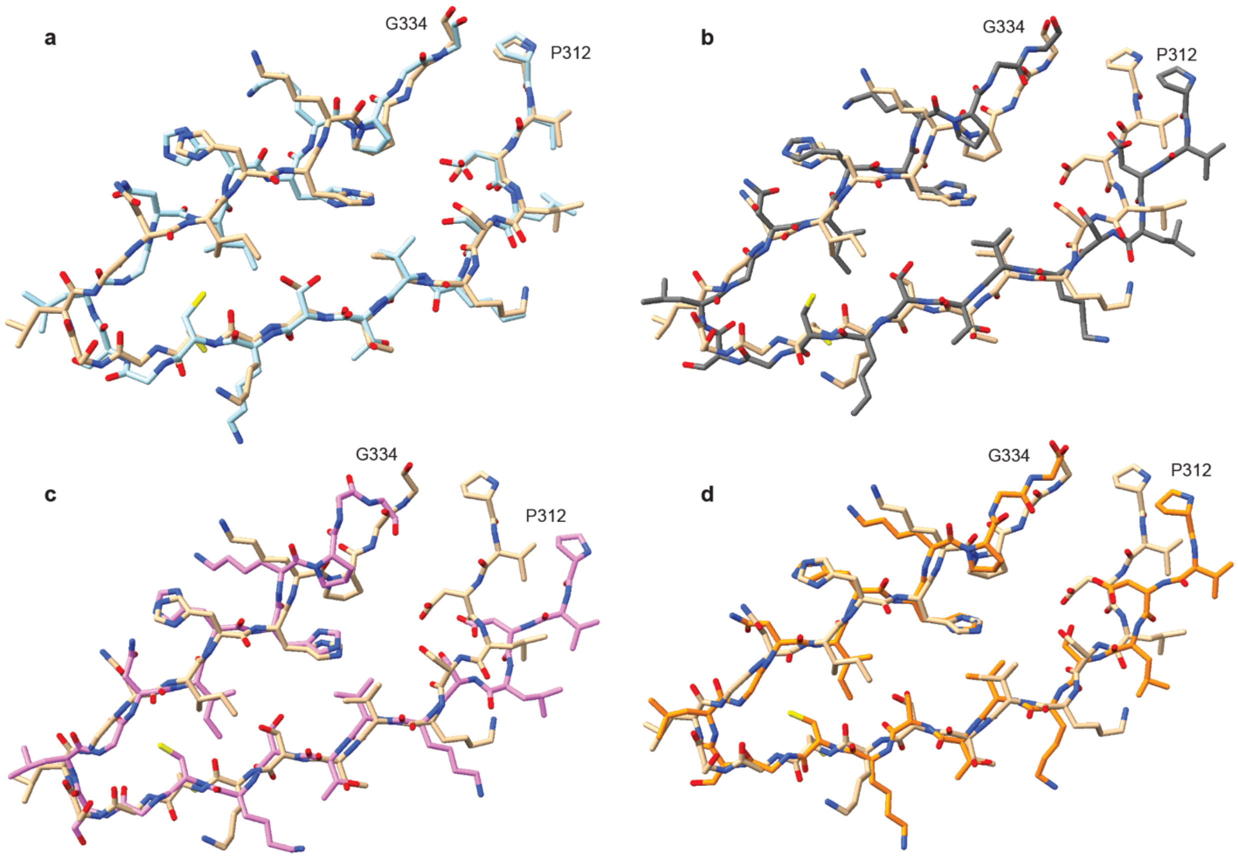

**Extended Data Fig. 4. Structural comparison between the R3 hairpin of the S305I Type I fold to R3 hairpins in other human tau folds.** (**a**), S305I Type II fold. (**b**), Sporadic AGD Type I fold (PDB: 7P6D). (**c**), P301T Type I fold (PDB: 9GG1). (**d**), CBD fold (PDB: 6TJO).

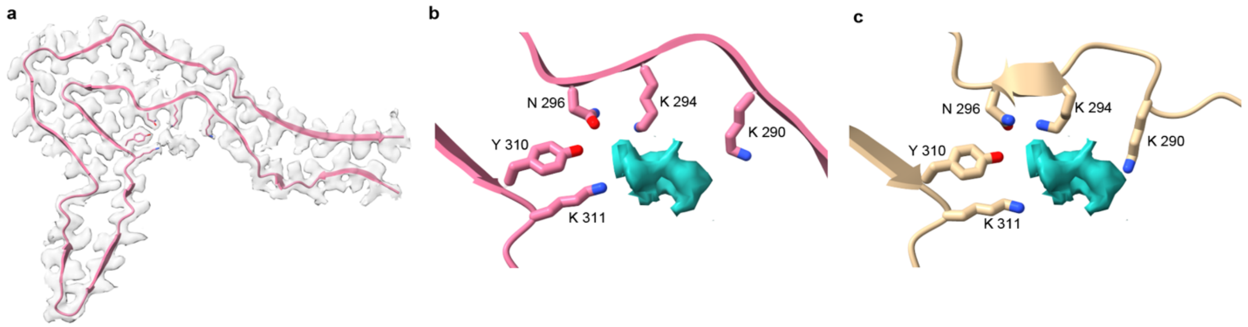

**Extended Data Fig. 5. P301T residues coordinate additional density in a similar motif to S305I.** (**a**), Map/model overlay of the P301T fold at threshold 0.00499 ($\sigma$ = 2.5) showing additional density in the same groove at that of S305I. (**b**), P301T residues coordinating additional density shown in blue. (**c**), The same residues in the S305I fold, with P301T additional density shown, demonstrating a very similar coordinating motif.

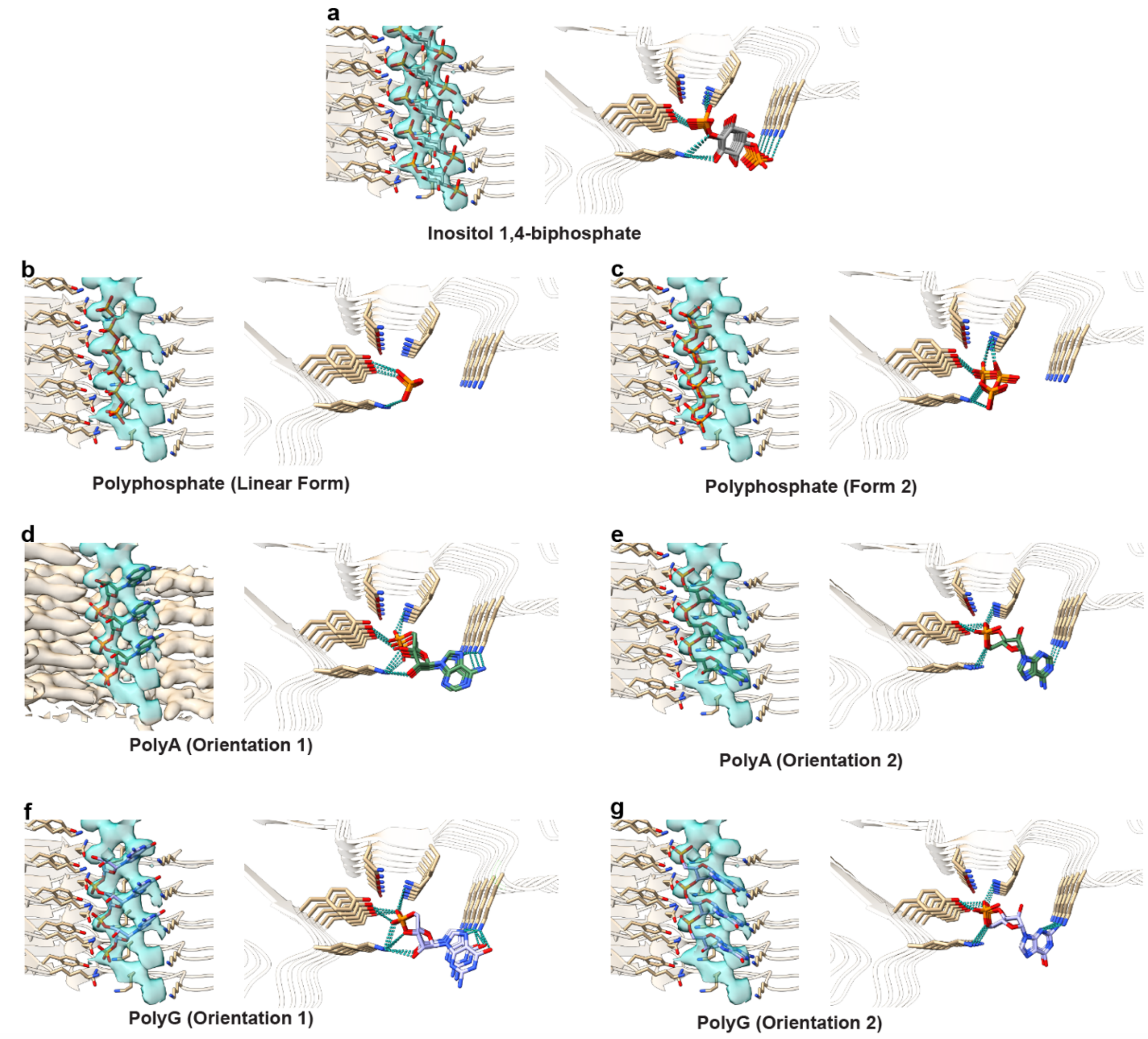

**Extended Data Fig. 6. Hypothesized cofactors that fit in the unassigned density of the S305I folds.** (**a**), Inositol 1,4-biphosphate. (**b**), Linear polyphosphate. (**c**), Form 2 polyphosphate. (**d**), PolyA RNA. (**e**), PolyA RNA in the opposite orientation from (**e**). (**f**), PolyG RNA. (**g**), PolyG RNA in the opposite orientation from (**f**).

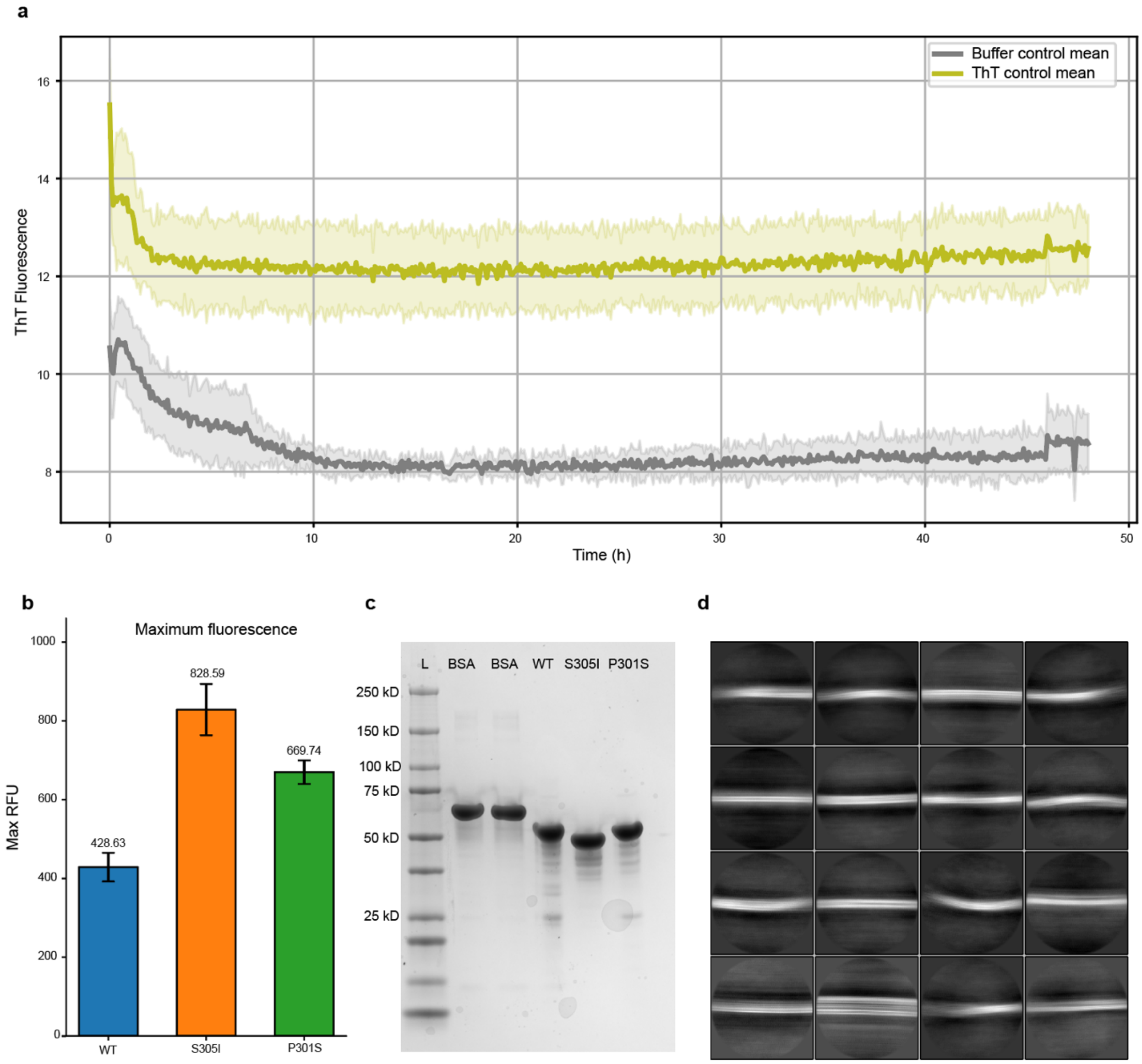

**Extended Data Fig. 7. Fibrilization assay controls.** (**a**), Fibrilization curves with ThT and buffer (top, yellow) and with buffer only (bottom, grey), showing that both ThT and protein are required for fluorescence signal. (**b**), Maximum fluorescence values for each tau construct. (**c**), Gel of WT, S305I, and P301S tau constructs, with BSA as a control. In each lane, 2 µg of protein were loaded. (**d**), 2-dimensional class averages of the S305I filaments represented in **Fig. 6d**.

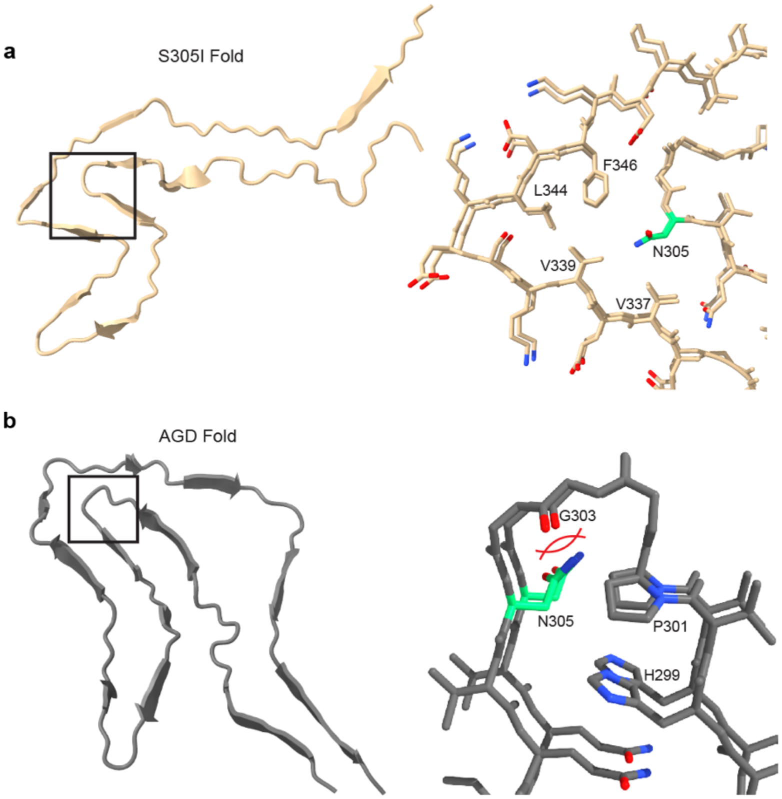

**Extended Data Fig. 8. S305N modeled into the S305I and AGD folds.** (**a**), N305 within the S305I Type I fold. The polar asparagine residue poorly complements the hydrophobic pocket in the S305I fold. (**b**), N305 within the sporadic AGD fold. N305 is not sterically accommodated in the pocket, clashing with either G303 (shown above) or H299, depending on the sidechain orientation.

**Extended Data Table 1. Cryo-EM data collection, refinement and validation statistics**

| | S305I Type I (EMDB-75195) (PDB 10IJ) | S305I Type II (EMDB-75196) (PDB 10IK) |
|---|---|---|
| **Data collection and processing** | | |
| Magnification | 105,000 | 105,000 |
| Voltage (kV) | 300 | 300 |
| Electron exposure (e–/Å$^2$) | 46 | 46 |
| Defocus range (μm) | -0.8 to -1.8 | -0.8 to -1.8 |
| Pixel size (Å) | 0.834 | 0.834 |
| Symmetry imposed | C1 | C1 |
| Initial particle images (no.) | 942,149 | 172,515 |
| Final particle images (no.) | 47,171 | 42,749 |
| Map resolution (Å) | 3.1 | 3.2 |
| FSC threshold | 0.143 | 0.143 |
| Map resolution range (Å) | 10-2.5 | 10-2.3 |
| **Refinement** | | |
| Initial model used (PDB code) | De novo | De novo |
| Model resolution (Å) | 3.4 | 3.5 |
| FSC threshold | 0.5 | 0.5 |
| Model resolution range (Å) | 18.1-2.16 | 29.4-2.19 |
| Map sharpening *B* factor (Å$^2$) | -63.4 | -58.3 |
| Model composition | | |
| Non-hydrogen atoms | 18,348 | 16,670 |
| Protein residues | 1,177 | 1,070 |
| Ligands | 0 | 0 |
| *B* factors (Å$^2$) | | |
| Protein | 74.57 | 89.13 |
| Ligand | N/A | N/A |
| R.m.s. deviations | | |
| Bond lengths (Å) | 0.004 | 0.006 |
| Bond angles (°) | 0.809 | 1.264 |
| Validation | | |
| MolProbity score | 2.66 | 2.88 |
| Clashscore | 12.61 | 15.99 |
| Poor rotamers (%) | 4.30 | 4.30 |
| Ramachandran plot | | |
| Favored (%) | 89.52 | 82.86 |
| Allowed (%) | 10.48 | 16.19 |
| Disallowed (%) | 0.00 | 0.95 |

**Extended Data Table 2. Localized RMSD comparisons between S305I Type I and other human tauopathy tau folds.**

| Human Tauopathy | PDB Code | Localized RMSD (Å) |
|---|---|---|
| S305I Type II | XXX | 2.80 |
| P301T Type I | 9GG1 | 5.04 |
| CBD Type I | 6TJO | 6.98 |
| AGD Type I | 7P6D | 7.50 |
| P301T Type II | 9GG6 | 8.29 |
| PiD | 6GX5 | 8.48 |
| ∆K281 PiD | 8P34 | 8.75 |
| CTE | 6NWP | 9.63 |
| SSPE | 8CAQ | 9.63 |
| AD SF | 5O3T | 10.08 |
| AD PHF | 5O3L | 10.19 |
| P301L | 9GG0 | 10.34 |
| V337M SF | 9EO9 | 10.45 |
| R406W SF | 9EOG | 10.59 |
| V337M PHF | 9EO7 | 10.59 |
| PSP | 7P65 | 10.70 |
| GPT | 7P6A | 10.73 |
| GGT | 7P66 | 10.86 |
